# Ultra-Sensitive Detection of Transposon Insertions Across Multiple Families by Transposable Element Display Sequencing

**DOI:** 10.1101/2024.08.21.608910

**Authors:** Pol Vendrell-Mir, Basile Leduque, Leandro Quadrana

**Affiliations:** Institute of Plant Sciences Paris-Saclay, Centre Nationale de la Recherche Scientifique, Institut National de la Recherche Agronomique, Université Evry, Université Paris-Saclay, 91405 Orsay, France

**Author notes:** Both authors contributed equally (Pol Vendrell-Mir, Basile Leduque).

## Abstract

**Background:** Mobilization of transposable elements (TEs) can generate large effect mutations. However, because new TE insertions are challenging to detect and transposition is typically rare, the actual rate and landscape of new insertions remains unexplored for most TEs.

**Results:** Here, we introduce a TE display sequencing approach that leverages target amplification of TE extremities to detect non-reference TE insertions with high sensitivity and specificity. By implementing this approach on serial dilutions of genomic DNA from *A. thaliana* lines carrying different repertoires of new TE insertions, we show that the method can detect TE insertions that are present at frequencies as low as 1:250 000 within a DNA sample. In addition, TE display sequencing can be multiplexed to simultaneously detect insertions for distinct TE families, including both retrotransposons and DNA transposons, increasing its versatility and cost-effectiveness to investigate complex “mobilomes”. Importantly, when combined with nanopore sequencing, this approach enables the identification of insertions using long-reads and achieves a turn around time from DNA extraction to insertion identification of less than 24h, significantly reducing the time-to-answer. Analysis of TE insertions in large populations of *A. thaliana* plants undergoing a transposition burst demonstrate the power of the multiplex TE display sequencing to assess the rates and allele frequencies of heritable insertions, enabling its implementation to study large-scale ‘evolve and resequence’ experiments. Furthermore, we found that ∼6% of de novo TE insertions show recurrent allele frequency changes consistent with either positive or negative selection.

**Conclusions:** TE display sequencing is an ultra-sensitive, specific, quick, and cost-effective approach to investigate the rate and landscape of new insertions for multiple TEs in large scale population experiments. We provide a step-by-step experimental protocol as well as ready-to-use bioinformatic pipelines, ensuring straightforward implementation of the method.

## INTRODUCTION

Transposable elements (TEs) are sequences that can move and replicate around the genome. TEs are categorized in two broad classes based on their mobilization mechanisms. Class I retrotransposons move via an RNA intermediate, while Class II DNA transposons use a “cut and paste” mechanism instead. TEs are further divided into superfamilies and families based on particular sequence features, such as the presence of specific terminal repeats or conserved protein domains [1].

Although TEs have long been viewed as mere genomic parasites [2], this view has dramatically shifted and it is now clear that TEs are key players in the functional as well as the structural evolution of genomes. Notwithstanding, the extent to which TE insertions contribute to heritable phenotypic variation remains unknown in most eukaryotes. This lack of knowledge is largely due to their repetitive nature, which makes TEs less amenable than single copy DNA to genome-wide analyses, particularly using short-read sequencing technologies. Moreover, TE mobilization is typically rare and therefore new TE insertions tend to be missed in small-scale population studies. Hence, being capable of detecting new TE insertions in large populations of individuals is essential to study transposition activity and its impact on genome function and evolution.

Several methods have been developed to detect new TE insertions using short-or long-read sequencing technologies. These methods rely on the selective sequencing of TE extremities, either using capture approaches [3–5], target DNA cleavage [6], or specific amplification [7–10] for library preparation. Capture approaches rely on the use of biotinylated oligos to selectively pull-down DNA fragments with complementary sequences [3–5]. Although these methods provide good sensitivity, designing specific oligos is challenging and their production costly, limiting their wide implementation. Approaches combining Cas9-mediated cleavage and adaptor ligation followed by long-read sequencing has been recently proposed as an interesting alternative, as it allows to sequence native DNA without additional steps [6]. However, the enrichment efficiency of such methods is limited, precluding its use for population level studies. Conversely, approaches based on target amplification of specific sequences theoretically offer the highest sensitivity. These methods have primarily been used to detect somatic insertions, but their applicability in investigating germline insertions remains to be determined. Additionally, PCR amplification can generate chimeric or non-specific products, requiring case-by-case modifications of experimental and bioinformatic methods to remove false positives. In fact, methods based on target amplification have been tailored to study specific TEs, mostly mammalian L1s or ERVs, which raises questions about their versatility and applicability to analyze diverse types of TEs, including both retrotransposons and DNA transposons.

Here, we describe a cost-effective (less than 12 EUR per sample’s library), simple, and ultra-sensitive assay to detect TE insertions in genomes compatible with short as well as long-read sequencing techniques. The method is based on classical low-throughput transposon display assays [7,11], but includes several modifications that make it readily compatible with high-throughput sequencing. We demonstrate its utility to investigate germline insertions using the model plant *Arabidopsis thaliana*, and we show that the method provides ∼250,000-fold enrichment of target TEs, enabling the detection of low-frequency insertions in large populations. In addition, we show that the obtained libraries can be analyzed using nanopore sequencing technologies, leading to longer reads and, importantly, a turn around time from DNA extraction to insertion identification of less than 24h, significantly reducing the time-to-answer. Last, we provide a detailed hands-on protocol (https://www.protocols.io/view/ted-seq-c7seznbe), including critical steps and quality controls, as well as a bioinformatic pipeline to analyze the obtained sequencing results (TEd-seq-pipe).

## RESULTS

### TE display sequencing

Classic transposon display assays rely on genomic DNA digestion using restriction enzymes [7,11] followed by ligation of a custom adapter and PCR amplification using one primer specific to the target TE sequence and another complementary to the adapter. To adapt this assay for high-throughput TE insertion detection using Illumina NGS, we replaced enzymatic digestion with random fragmentation by sonication, producing fragments of approximately 200-500 bp (Fig 1a). Similar to standard NGS library preparations, the fragmented DNA is end-repaired and A-tailed to facilitate the ligation of custom adapters, which are compatible with Illumina sequencing. These adapters were designed to exclude amplification by adapter primers only, avoiding the generation of random whole genomic library. Specifically, the priming sites are not present on the adapter *ab initio*, but generated through a first round of extension from the TE primer, analogous to the vectorette/splinkerette PCR strategies [12,13]. Unlike these strategies, which utilize hairpins or partially-mismatched “bubbled” adapters, the present method relies on asymmetric Illumina forked-adapters. These adapters are much shorter and readily compatible with NGS libraries, providing advantages compared to vectorette/splinkerette PCR adapters. To block extension from the asymmetric forked-adapter, its short arm includes a 3’ dideoxy nucleotide (Fig 1b). Even in the unlikely situation where an adapter is extended first, producing templates with full adapter sequences at both ends, these fragment ends will preferentially anneal to form an intramolecular “panhandle” structure, which cannot be amplified further. The “suppression PCR” effect occurs because the complementarity of the adapter’s primer is shorter than that of the adapter itself, favoring intramolecular pairing. In combination, the universal adapter’s design and primer sequences ensure that only *bona fide* PCR templates containing both the target TE site and adapter sequence are generated during the first rounds of extension.

**Figure 1.**
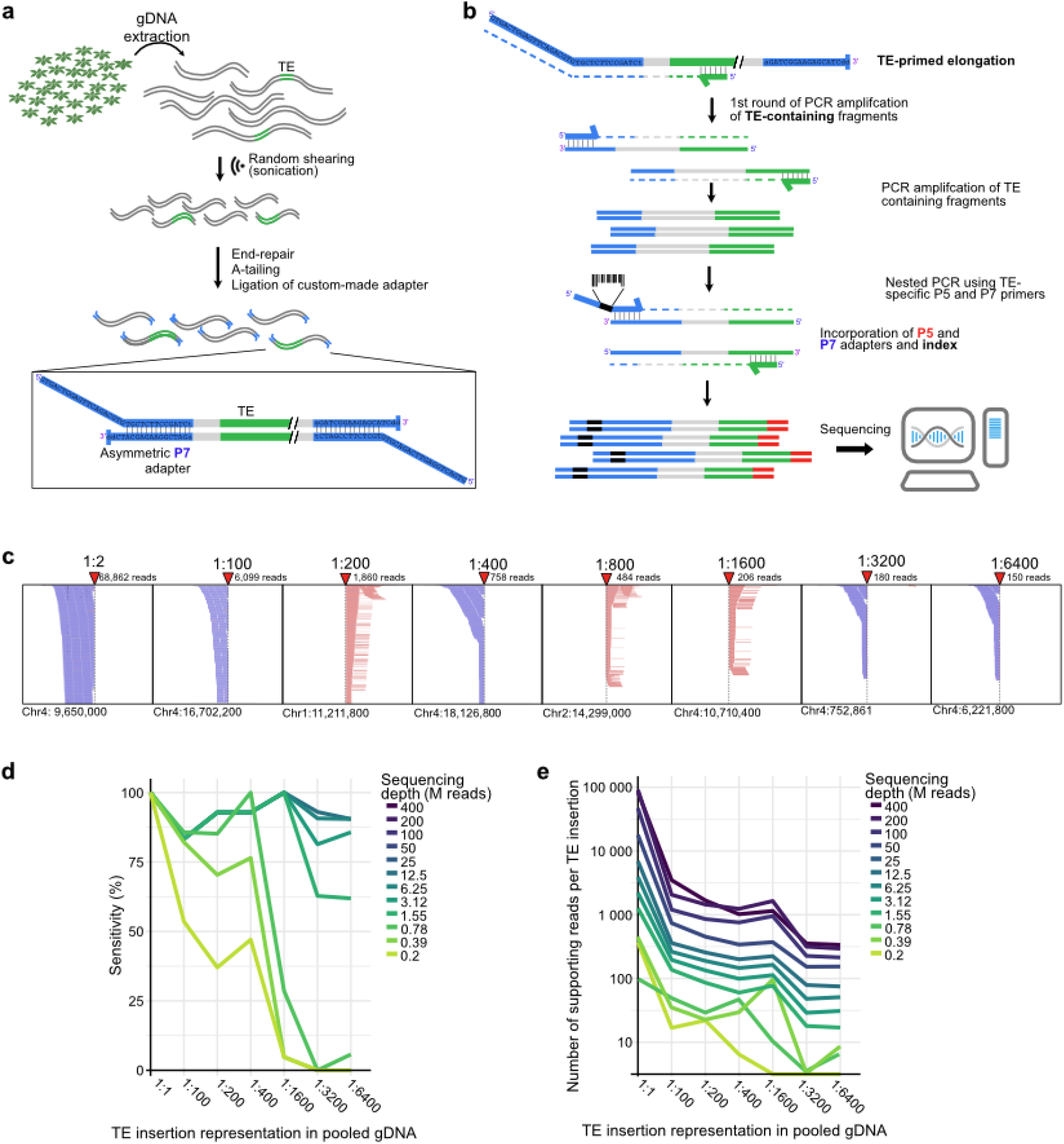
Transposable display sequencing method. a. Schematic representation of the initial steps to prepare adapter-containing whole genome libraries. **b**. Schematic representation of specific amplification of TE-containing adapted fragments and incorporation of P5 and P7 adapters compatible with illumina sequencing. **c**. Examples of TE insertions detected at different dilution factors. **d**. Sensitivity of TEd-seq to detect TE insertions at different dilution factors and in relation to sequencing variable depth. **e**. Number of reads supporting each TE insertion present at different dilution factors in relation to variable sequencing depth.

To increase further the sensitivity and specificity of the assay, we included a nested PCR step using a TE-specific primer closer to the TE edge together with an adapter’s primer (Fig 1b), respectively containing the P5 and P7 sequences required for cluster recognition on Illumina sequencing machines. The adapter’s primers also include sequence barcodes, enabling the multiplexing of up to hundreds of samples in the same sequencing run. Importantly, to improve sequencing quality due to the low-diversity of the amplicon library, nested TE-specific primers contain different numbers of spacer nucleotides before the P5 adapter. Such simple modifications significantly improve library diversity and overall sequencing quality.

After pair-end sequencing and demultiplexing, reads are processed bioinformatically to eliminate PCR duplicates and select paired-reads containing the target TE sequence end, which is trimmed to retrieved informative insertion sites sequences. Alignment on the reference genome assembly and clustering of reads therefore allows the detection of reference and non-reference insertion sites (Fig 1c).

### Highly specific and ultra-sensitive detection of new TE insertions

To assess the sensitivity and specificity of the Transposable Element display sequencing (TEd-seq) to detect TE insertions, we took advantage of an epigenetically-compromised population of *Arabidopsis thaliana* lines, the so called epiRILs, that have underwent extensive transposition during six generations of selfing. The identity and genomic location of heritable TE insertions in each of these lines were previously characterized using Illumina mate-pair libraries [5,14] and TE sequence capture [5], providing a gold standard data set to test the sensitivity and specificity of TEd-seq. We selected eight lines carrying distinct repertoires of non-reference TE insertions of the LTR-retrotransposon *ATCOPIA93* (also known as *Évadé*). We extracted genomic DNA (gDNA) from each of these epiRILs and serially diluted them at different concentrations (from 1:50 to 1:6400). Pooling together these differentially diluted gDNA thus leads to a mixed gDNA with TE insertions present at different dilutions. The resulting pooled DNA was subjected to TEd-seq using primers specific to the mobile *ATCOPIA93* copy (*AT5TE20395*), sequenced to a depth of 400M pair-end reads, and analyzed bioinformatically to detect non-reference TE insertion sites. Comparison between the set of non-reference *ATCOPIA93* insertions detected by TEd-seq to the gold standard set of previously identified insertions [14] showed 100% correspondence for all the dilutions, demonstrating the high specificity of this enrichment approach. In addition, 87-100% of TE insertions were detected for all the dilutions (Fig 1c, d). Importantly, the number of reads supporting each *ATCOPIA93* insertion correlated well with the dilution factor (Fig 1c,e), indicating that the number of supporting reads is a reliable estimator of the TE insertion representation.

To determine the sensitivity and specificity of TEd-seq for variable depth of sequencing, we subsampled the number of row paired-end reads obtained in the TEd-seq experiment and detected non-reference TE insertions. As expected, sensitivity decreased in relation to sequencing depth (Fig 1d). Specifically, 80% of the TE insertions present in the 1:6400 dilution were detected with as little as 6M pair-end reads. Based on this result, we estimate that 400M reads is sufficient to detect insertions presented in as low as 1:250,000 dilution.

### Detecting TE insertions using ONT sequencing

Recent developments in sequencing technologies based on Oxford Nanopore Technologies (ONT) are revolutionizing the output and speed of genomic analysis. We reasoned that in addition to allowing the sequencing of long DNA fragments, which can facilitate the identification of insertion sites within highly repetitive or low complexity regions, its portability and real-time generation of sequencing data can significantly reduce the time-to-answer. We thus tested the possibility of sequencing a TEd-seq library using ONT (Fig 2b). We optimized the fragmentation and size-selection steps to obtain longer DNA fragments, which are more suitable for ONT sequencing. In addition, because ONT sequencing is not affected by library complexity, TE-specific primers do not include spacer nucleotides.

**Figure 2.**
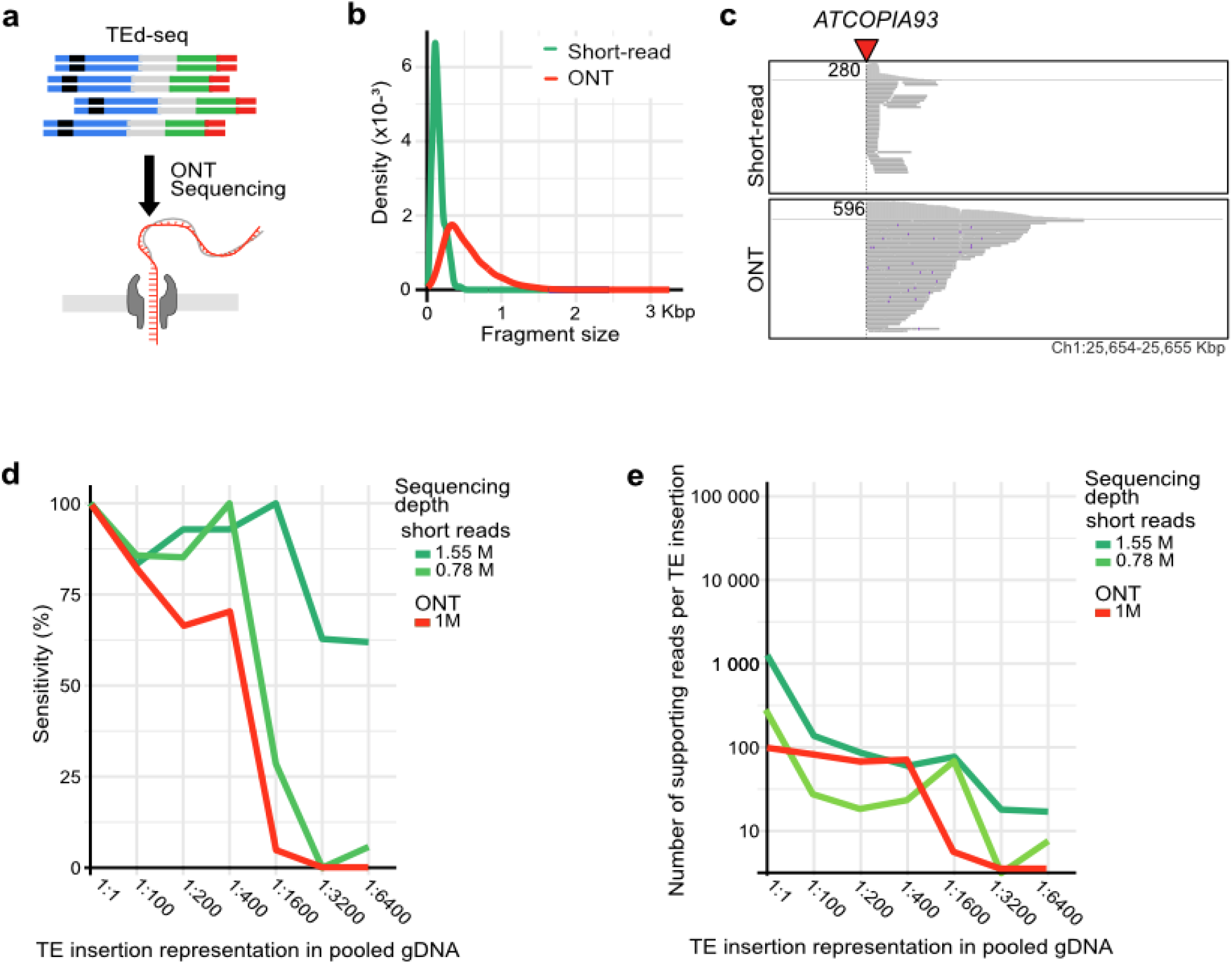
Transposable elements display long-read sequencing. Schematic representation of TEd-seq sequencing using ONT. b. Fragment size distribution of short-read and ONT-based TEd-seq. C. Example of a TE insertion detected by short-read and ONT TEd-seq. **d**. Sensitivity of ONT TEd-seq to detect TE insertions at different dilution factors. **e**. Number of reads supporting each TE insertion present at different dilution factors for both short-read and ONT TEd-seq.

We performed a TEd-seq library of the retrotransposon *ATCOPIA93* using gDNA extracted from a single epiRIL line (epiRIL500) diluted in 1:200 in Col-0 gDNA. As expected, the average size of ONT reads is longer than the fragments obtained by NGS (814.27 vs 350 nt, respectively), with some reads reaching more than 3000 nt (Fig. 2b and c). We next detected non-reference TE insertions using a modified bioinformatic pipeline compatible with ONT reads. Comparison between the set of non-reference *ATCOPIA93* insertions detected by TEd-seq to the gold standard set of previously identified insertions [14] in this epiRIL showed 100% correspondence, demonstrating that combination of TEd-seq with long-read ONT sequencing is highly specific. To test the sensitivity of ONT sequencing to detect non-reference TE insertions, we generated and sequenced the TEd-seq library containing the mixed pool of serially diluted gDNA from the different epiRILs. With as little as 1M ONT reads we identified more than 70% of insertions present in 1:400 dilution (Fig. 2d). This sensitivity is close to the one obtained by ∼1M short-reads, indicating that the major factor affecting the sensitivity of TEd-seq is the depth of coverage. In combination, these results establish that ONT can be used to sequence TEd libraries. Importantly, the portability and short hand-off time of ONT sequencing offers a turn around time from DNA extraction to insertion identification of less than 24h, expediting the investigation of TE mobilization for routine experiments.

### Simultaneous detection of insertions from multiple TE families using multiplex TEd-seq

Eukaryotic genomes typically contain several active TE families. Thus, to effectively investigate transposition activity, it is desirable to simultaneously detect non-reference insertions for multiple TE families. To explore this possibility, we designed TEd-seq primers specific for the CACTA and MuDR DNA transposons *ATENSPM3* and *VANDAL21*, which are highly mobile in the epiRILs population [14]. Using gDNA from epiRILs 454 (dilution 1:10) and 500 (dilution 1:200) we generated, for each epiRIL, independent TEd-seq libraries for *ATCOPIA93, VANDAL21* and *ATENSPM3*, together with multiplexed TEd-seq libraries generated using an equimolar mix of the three TE-specific primers during the amplification steps (Fig. 3a). Based on 1M ONT reads, we detected insertions for the three TE families (Fig. 3b). Remarkably, all homozygous non-reference insertions previously identified in the epiRILs [14] were detected in our multiplexed TEd-seq experiments (Fig. 3c,d), demonstrating a 100% sensitivity and the possibility to study several TE families simultaneously. Importantly, the multiplexed TEd-seq libraries yielded better sensitivities than TE-specific libraries (Fig. 3b), possibly due to higher sequence diversity of multiplexed libraries. In addition, out of the 96 non-reference TE insertions detected in the multiplexed libraries, only three appear to be false positives, leading to an overall precision of 96.8% (Fig. 3c,d). Altogether, these results established multiplex TEd-seq as a remarkably cost-effective, highly sensitive, and specific method to simultaneously detect non-reference insertions of different types of TEs, enabling to investigate transposition activity of rich mobilomes.

**Figure 3.**
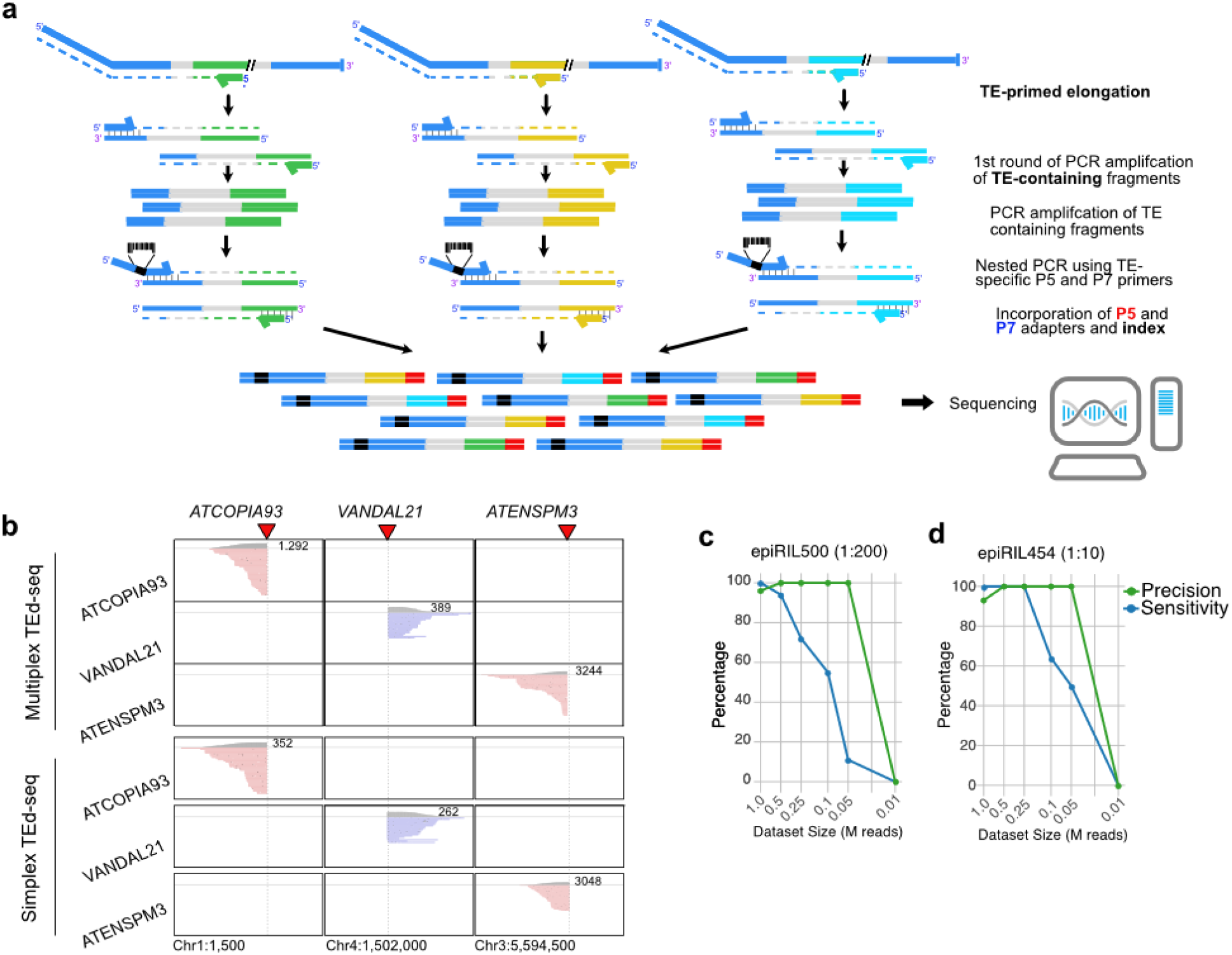
Simultaneous detection of insertions for different families by multiplexed transposable element display sequencing. a. Schematic representation of specific multiplexed amplification of TE-containing adapted fragments and incorporation of P5 and P7 adapters compatible with illumina sequencing. **b**. Examples of TE insertions from different TE families detected by multiplex and simplex TEd-seq. **c** and **d**. Precision and sensitivity of multiplex TEd-seq to detect TE insertions present in different epiRILs.

### Multiplex TEd-seq empowers evolve and resequencing experiments

Although most TE-induced mutations are likely deleterious, empirical and theoretical evidence indicates that a continuum of selective effects must exist (29). However, empirical estimation of the evolutionary dynamics of new TE insertions remains largely underexplored. Whole genome sequencing (WGS) of pools of individuals from experimentally evolved populations, such as in common garden competition experiments or experimental evolution setups, has enabled the identification of genomic responses to selection. The so-called “evolve and resequencing” (E&R) approaches use short-read Illumina WGS to identify segregating polymorphisms and characterize their changes in allele frequency across generations. Detection of complex DNA sequence variations, such as TE insertion polymorphisms, as well as of rare variants remains very challenging using WGS, limiting the implementation of E&R approaches to investigate the evolutionary dynamics of new TE insertions. The ultra-high sensitivity of TEd-seq combined with its ability to simultaneously detect non-reference TE insertions for multiple TE families in a cost-effective manner may therefore empower E&R experiments. As a proof-of-concept of the utility of TEd-seq to quantify the presence and frequency of non-reference TE insertions in E&R experiments, we generated a largely isogenic wild-type population of Arabidopsis plants undergoing a burst of transposition for the endogenous TE families *ATCOPIA93, VANDAL21*, and *ATENSPM3*. Specifically, we crossed wild-type Col-0 plants with distinct epiRILs (epiRIL523, 54, and 393) containing active copies for these three TEs families. F1 plants were selfed to obtain thousands of F2 seeds. An equal number of F2 seeds (3500) from each cross, together with naïve Col-0 seeds (500), were mixed to obtain a so-called “TE invaded” population (G0) (Fig 4a). We performed two parallel E&R experiments by growing 2500 plants from the TE invaded population in environmental chambers reproducing realistic climates, including temperature, humidity, and light intensity and quality (Supplementary Figure 1) recorded at an experimental station near Paris (see methods). All produced seeds at the end of the life cycle were collected from each population. This was repeated for another generation to obtain G2 offspring.

**Figure 4.**
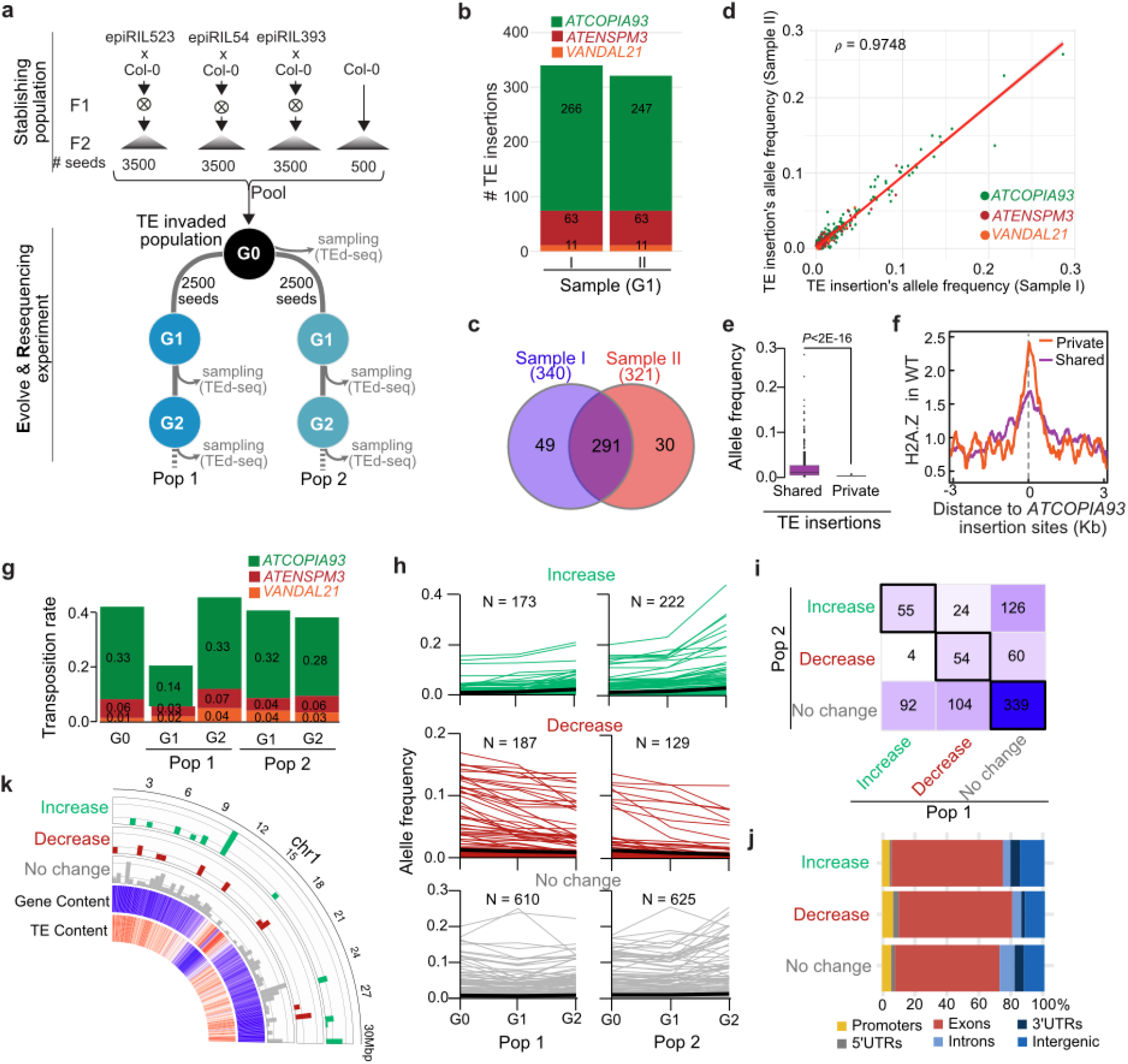
Investigation of the evolutionary dynamics of new TE insertions using TEd-seq. a. Schematic representation of the evolve and resequencing (E&R) experiment developed to study the evolutionary dynamics of new TE insertions. Two independent populations (Pop. 1 and Pop. 2) were used to run two E&R replicated experiments. B. Number of TE insertions identified by multiplex TEd-seq in two independent samples of 1000 offspring of G1 Pop. 1 for the three TE families (*ATCOPIA93, VANDAL21*, and *ATENSPM3*) transposing actively in the experimental population. **c**. Number of private and shared insertions detected in the two samples. **d**. Comparison of allele frequencies of all TE insertions estimated in the two populations. **e**. Allele frequencies of shared and private TE insertions. Statistically significant differences are indicated (MWU test). **f**. Meta-analysis of levels of H2A.Z levels around shared and private *ATCOPIA93* insertion sites. **g**. Average number of private and shared insertions detected across generations in both replicated populations. **h**. Allele frequencies change across generations in both populations. Insertions showing increase, decrease, or no change allele frequencies were selected by comparison with allele frequencies trajectories calculated under genetic drift. **i**. Comparison of TE insertions detected as increase, decrease, or no change among populations. **j**. Fraction of TE insertions identified as recurrent increase, decrease, or no change within genic features and intergenic regions. **k**. Circos representation of TE insertions identified as recurrent increase, decrease, or no change in chromosome 1 of Arabidopsis. Density of annotated genes and TEs are indicated as heatmaps.

To assess the ability of TEd-seq to accurately estimate the allele frequency of non-reference TE insertions, we performed multiplex TEd-seq (*ATCOPIA93, VANDAL21*, and *ATENSPM3)* on two random samples of 1000 G1 offsprings from population 1. In total, we identified 340 and 321 non-reference insertions in each sample, most of which belonged to the LTR retrotransposon *ATCOPIA93* and were shared between samples (Fig 4b and c). We next estimated the allele frequency of each TE insertion by comparing the numbers of reads supporting new (i.e. non-reference) TE insertions in relation to that for fixed (i.e. reference) insertions. Remarkably, allele frequencies were highly consistent between samples (Fig. 4d), demonstrating that TEd-seq provides robust allele frequencies estimations of segregating TE insertions.

Shared insertions among offspring samples should correspond to segregating alleles in the G1, and as such they are expected to be present at higher allele frequencies than private insertions. To test this expectation, we compared the allele frequency of shared and private non-reference TE insertions. As predicted, shared insertions were covered by a much higher number of reads than private ones (Supplementary Figure 2), and we estimated that segregate at frequencies higher than 0.025, compared to 0.004 for the latter (Figure 4e). Thus, private insertions represent *de novo* heritable transposition events.

It was previously shown that *ATCOPIA93* integrates preferentially within nucleosomal DNA containing the histone variant H2A.Z [14]. Analysis of the integration sites of *ATCOPIA93* in relation to the presence of H2A.Z confirmed this preference, and revealed higher occupancy of H2A.Z at de novo than shared insertion sites (Figure 4f). This result may be suggesting counterselection of Te insertions within H2A.Z containing regions.

Next, we set out to investigate the transgenerational dynamics of active transposition on the E&R experiment. We performed multiplex TEd-seq on DNA extracted from samples of 1000 offspring each of G0, as well as G1 and G2 generations for both replicated populations (Fig. 4a). In total we identified 5799 insertions, 48% were private to individual samples. Using the number of private insertions as a proxy for de novo heritable transposition, we estimated the transposition rate across replicates and generations. Estimated transposition rates were very similar between populations and generations (Figure 4g), consistent with lineal accumulation of new TE copies across generations.

To characterize the evolutionary dynamics of segregating TE insertions, we estimated the allele frequency across generations of TE insertions shared among samples. To detect insertions potentially subjected to selection, we compared their allele frequencies with neutral expectations based on random genetic drift and considering less than 5% of outcrossing rate (ref). Most (∼63%) insertions segregate in random genetic drift fashion. Conversely, 20% and 16% of the TEs insertions show a statistically significant increase or decrease in allele frequencies, respectively. Moreover, 55 and 54 of these insertions show paralleled allele frequency changes in both populations (Fig. 4i). Such parallel changes are unlikely to be produced by chance (hypergeometric test p-value < 8.7E-5 and 1.7E-11 for increases and decreases, respectively), suggesting recurrent positive or negative selection of these TE insertions across replicated populations. In addition 339 TE insertions show no changes in allele frequency across populations, suggesting that they lead to neutral or nearly neutral mutations (Fig. 4i). The distribution of TE insertions with reproducible increase, decrease, or no changes in allele frequencies reveal no chromosome-wide bias (Figure 4k), suggesting that the observed changes in allele frequencies are not due to selection on underlying haplotypes (i.e. background selection) but rather on individual TE insertion. In addition, TE insertions with recurrent decrease in allele frequency show overrepresentation over exonic features compared to neutral TE insertion mutations (Figure 4j), consistent with this exonic insertions being more deleterious. Among the 42 genes with insertions in their gene body or promoter that are increasing in population frequency, we identified 20 enriched GO terms. These terms are primarily associated with various binding activities (e.g., nucleotides and nucleosides), transferases, and transmembrane transporters. This group of genes is also enriched at the cell periphery and plasma membrane, suggesting roles in cell-environment interaction and transport processes. Conversely, among the 45 insertions located within or near genes that are recurrently counterselected, we identified four enriched GO terms (GO:0045229, GO:007155, GO:0071554, and GO:0008194). These terms are mainly involved in cell wall structure and organization, as well as enzymatic activities related to glucosylation, which are essential for maintaining cell integrity and ensuring proper cellular interactions and responses to environmental changes.. Specifically, we found a *ATCOPIA93* and *ATENSPM3* insertion with consistent decrease in allele frequency across populations within the genes encoding FORMIN HOMOLOGY 5 (atFH5) and KNUCKLES (KNU), respectively involved in endosperm cellularization [15] and flower determinacy [16] (Supplementary Figure 3). In combination, these results indicate that as much as 6% of new TE insertions are affected by positive or negative selection in our E&R experiment, and demonstrate the power of our combined experimental and analytical approach to investigate experimentally the evolutionary dynamics and impact of new TE insertions.

## DISCUSSION

Here, we adapted the classical Transposable Element display method to be readily compatible with modern sequencing technologies. We demonstrate that such an approach provides ultra-sensitive and specific detection of non-reference TE insertions, enabling high-throughput identification of very rare insertions in a large population of individuals. We have optimized TEd-seq to be able to detect insertions for several TE families simultaneously, including the LTR-retrotransposon *ATCOPIA93*, and the DNA transposons *VANDAL21* and *ATENSPM3*. Such a multiplex TEd-seq approach offers a powerful approach to study complex mobilomes, such as the one of the model plant Arabidopsis thaliana. Its simplicity and versatility significantly facilitate its implementation to study diverse TE families, in this as well as in other species (reference to PIF/Harbinger paper).

Our results demonstrate that TEd-seq is compatible with both short-as well as long-read sequencing technologies. Compared to the short-read sequencing, TEd-seq with ONT produces reads that range between 0.5Kbp till 3Kbp, unlocking the detection of non-reference TE insertions within highly repetitive sequences, such as satellite repeats, telomeres and centromeres. Furthermore, we show that the sensitivity and specificity of TE insertion identification is comparable between both sequencing technologies. Importantly, the portability and little hand-off time of ONT sequencing offers a turn around time from DNA extraction to insertion identification of less than 24h, expediting the investigation of TE mobilization for routine experiments. Furthermore, we estimated that the per-sample price for a 24-plex library is about 12 USD, offering a cost-effective method. Its simplicity and low outfront cost, together with the steady decrease in the cost of ONT sequencing, will significantly help to study transposition across genomes.

The combination of ultra-high sensitivity, accuracy, and cost-effectiveness are particularly suitable for large scale experiments. As a proof-of-concept, we implemented multiplex TEd-seq to investigate transposition dynamics across generations in populations of Arabidopsis plants undergoing a transposition burst. Our results show that the global transposition rate is relatively constant across generations. This observation is at odds with the notion that once initiated, TE invasion follows a chain reaction as every new copy may serve as a source for new transposition events. Unlike previous experimental populations, which are akin to mutation accumulation lines (Quadrana 2019), our E&R approach involved thousands of individuals growing in a realistic natural environment. Under such circumstances, moderately deleterious mutations could be purged by natural selection, potentially limiting runaway accumulation of new TE insertions. Consistent with this scenario, ∼15% of segregating TE insertions show steady decrease in allele frequency in both population replicates, consistent with measurable counter selection. Given that our proof-of-concept E&R has only two generations, our estimation of counter selected TE insertions is likely underpowered. Extending this experiments to additional generations, as well their replication across distinct environments, would provide precise fitness effect estimations of new TE insertion mutations

## Conclusions

TEd-seq enables the simultaneous study of heritable transposition for several TEs and in a large number of individuals. Implementing TEd-seq on E&R experiments will empower the investigation of the evolutionary dynamics of new TE insertions in real time. We provide a step-by-step protocol and bioinformatic pipelines to readily implement TEd-seq to study ebay TE in any organism.

## Material and methods

### Plant material

The *A. thaliana* Col-0 and the epiRILs lines used in this work were described before [5,14]. To generate the TE invaded populations, three epiRILs (epiRIL54, epiRIL523 and epiRIL393) were crossed with Col-0 and an equal number of seeds from independent F2 segregating populations were collected and mixed together to generate a founder population. 1000 seeds of this mix were sowed and plants were grown for the whole life cycle in a growth chamber (Percival) under environmental conditions simulating the climate of February 2022 to June 2022 (30’ resolution) nearby Paris (48°16’55.1”N 2°40’25.3”E) (Supplementary Figure 1). All seeds were collected and pooled, germinated on 1/2MS media and grown under standard conditions. 10 days old seedlings were collected in bulk, flash frozen in liquid nitrogen, and stored at -80°C until use. Plant material was grinded using pestle and mortar, and used for DNA extraction with the CTAB method [17].

### TEd-seq protocol for short-reads

50 μl of 20 ng/μl gDNA was sonicated in a Covaris M220 sonicator (Peak incident power 75W, Duty factor (18.6%), 200 Cycles per burst, for 175 seconds) to obtain fragments of 200-600 bp. Fragmented DNA was subjected to end repair and A-tailing using the NEBNext Ultra II End Prep Enzyme Mix (NEB, Ref:E7645) in a total volume of 60 μl, followed by ligation using TED-seq forked-adapters (final concentration of 0.8 μM), NEBNext Ultra II Ligation Master Mix, and NEBNext Ligation Enhancer. Adapter-ligated DNA was size selected using AMPure XP Beads (Beckman Coulter, ref:A63880) and used as template for selective amplification of TE containing fragments with NEBNext Ultra II Q5 Master Mix PCR and primers TEDseq_outer_for and TE_outer_rev. The following PCR conditions were used: 98°C for 30 sec, followed by 98°C for 10 sec, 61°C for 75 sec for 20 cycles with a last cycle of 5’ at 61°C

PCR product was purified and size selected using AMPure XP Beads (0.9X). The DNA purified product was subjected to a second round of PCR with nested primers containing custom illumina P5 and barcoded P7 adapters (P7_Primer_index#, P5_TE_primer). The PCR conditions were: 98°C for 30 sec; 6 cycles of 98°C for 10 sec and 61°C for 75 sec; 10 cycles of 98°C for 10 sec and 72°C for 75 sec; and a final step at 72°C for 5 min. Finally, the DNA library was purified and size selected using AMPure XP Beads (0.9X) and resuspended in 30μl of TrisHcl 10mM. Short-read sequencing was performed by BGI Tech Solutions (Hong Kong). Multiplexed TED-seq was conducted using equimolar concentrations of TE_outer_rev specific to each TE family (TE_outer_rev_ATCOPIA93: GTGAGTCCTCTTCAACGGCT; TE_outer_rev_VANDAL21: GGTTCGGATGGTTTGGTTCGG; TE_outer_rev_ATENSPM3: CGAAGCTGTAGAGGGAAGTG) and P5_TE_primers (see supplementary Table 1).

### TEd-seq protocol for long-reads

50 μl of 20 ng/μl gDNA was sonicated using a Covaris sonicator (50W, 2% duty factor, 200 cycles per burst, 3 minutes) to obtain 750-1500 bp fragments. Fragmented DNA was processed with the TEd-seq protocols described above, adjusting the AMPure XP Beads molarity rate (0.7X) to select fragments larger than 700 bp. 1 μg of pooled TEDseq libraries was subjected to end repair and A-tailing using the NEBNext® FFPE DNA Library Prep Kit (NEB:E6650S). Nanopore adapter ligation and library construction was conducted using the Nanopore Ligation Sequencing Kit (ONT, SQK-LSK110) following manufacturers instructions. Libraries were loaded on a R9.4.1 flowcell (ONT) according to the manufacturer’s instructions. A standard 72 h sequencing was performed on a PromethION 2 Solo (ONT). EpiRIL454 TED-seq ONT library was conducted using the Nanopore Ligation Sequencing Kit (ONT, SQK-LSK114 and sequenced on a R10.4.1 flowcell (ONT). MinKNOW software (version 23.11.7) was used for sequence calling.

### TEd-seq short-reads bioinformatic pipeline

Short-read libraries were demultiplexed by TE family, and TE sequences were trimmed from both ends using cutadapt (V2.6)[18] with options: -e 0.1, -O 25. PCR duplicates were removed using bbmap clumpify (V38.18) with options: -subs=2, dedupe, -Xmx32g. Reads were mapped to the reference genome using bowtie2 (V2.3.5.1)[19] with parameters: --local, --very-sensitive. TIP insertions were called using a custom pipeline accessible in GitHub. Briefly, read coverage was estimated at each genome position; positions with more than 30 reads for *ATCOPIA93*, 10 reads for *ATENSPM3*, or 3 reads for VANDAL21 were considered as possible new insertions. Insertions overlapping reference TEs of the same family and in plastid genomes were excluded from the analysis. TE insertions detected in different samples and located less than 500bp were considered to be shared

### TEd-seq long-reads bioinformatic pipeline

Raw reads were basecalled using guppy basecaller (V6.4.6+ae70e8f) for R9 libraries and Dorado (V0.6.0) for R10 libraries. Reads were then trimmed and demultiplexed with cutadapt (V2.6)[18] using the barcoded custom P7 sequence and the P5 TE sequence with options: -e 0.2, -O 25. Reads containing multiple adapters were discarded. Trimmed and demultiplexed reads were mapped to the reference genome using minimap2 (V2.17-r941) [20]. Finally, an adapted TED-seq analysis pipeline was run to detect TIP germline insertions (accessible at the GitHub repository).

### Allele frequency and transposition rate estimations

Three independent samples of 1000 seeds from each population and from generations 1 and 2 were grown for 10 days in 1/2MS medium. DNA was then extracted in bulk from all the seedlings of each sample. These three independent samples served as biological replicates to estimate allele frequency. For generation 0, due to a limited number of seeds, only 1000 seeds were sown. Multiplexed short-read TEDseq was performed on each individual sample as described above. For generation 0, we created two independent short-read TEDseq libraries from the same DNA extraction and treated them as technical replicates.

To estimate allele frequencies for shared and non-shared TE insertions, the number of second-in-pair reads per sample at each insertion site were counted and normalized by the average number of second-in-pair reads across reference TE insertions. Insertions with average allele frequencies lower than 2‰ in G0 and G1 were discarded. Using the observed allele frequencies from Generation 0 (p), the expected frequencies were estimated assuming a mixing mating system with a selfing rate (s) of 0.95. Changes in allele frequency due to genetic drift were identified by comparing predicted to observed frequencies in each generation using a Z-test. Insertions with a p-value lower than 0.05, following the same trend across generations (increasing from G0 to G1 and G1 to G2, or decreasing in the same manner), were categorized as changing in frequency.

To estimate the transposition rate, the reads for each TE family from the E&R were demultiplexed and downsampled to the same number of reads using Seqtk subseq (V1.3-r106). TED-seq germline short-reads pipeline was then run. Windows of 1000 bp were opened from each TIP position, and each insertion was categorized as private or shared based on whether it was detected in a single sample or across multiple samples, respectively. Private insertions from each sample were considered for estimating the transposition rate.

## Acknowledgements

We thank members of the Quadrana group, Sayuri Tsukahara, and Tetsuji Kakutani for discussions and advice.

## Authors contributions

LQ conceived the project and wrote the manuscript. LQ and BL established the TEd-seq protocol and bioinformatic analysis, with additional input from PV. PV developed the multiplex and long-read TEd-seq, and performed population experiments and analyses. All authors interpreted the data, read, and approved the paper.

## Data and material availability

Original scripts are available on GitHub. Sequencing data are available at the European Nucleotide Archive (ENA) under project PRJEBXXXX. A detailed TEd-seq method is available at *https://www.protocols.io/view/ted-seq-c7seznbe*

---

**Supplementary Figure 1.**
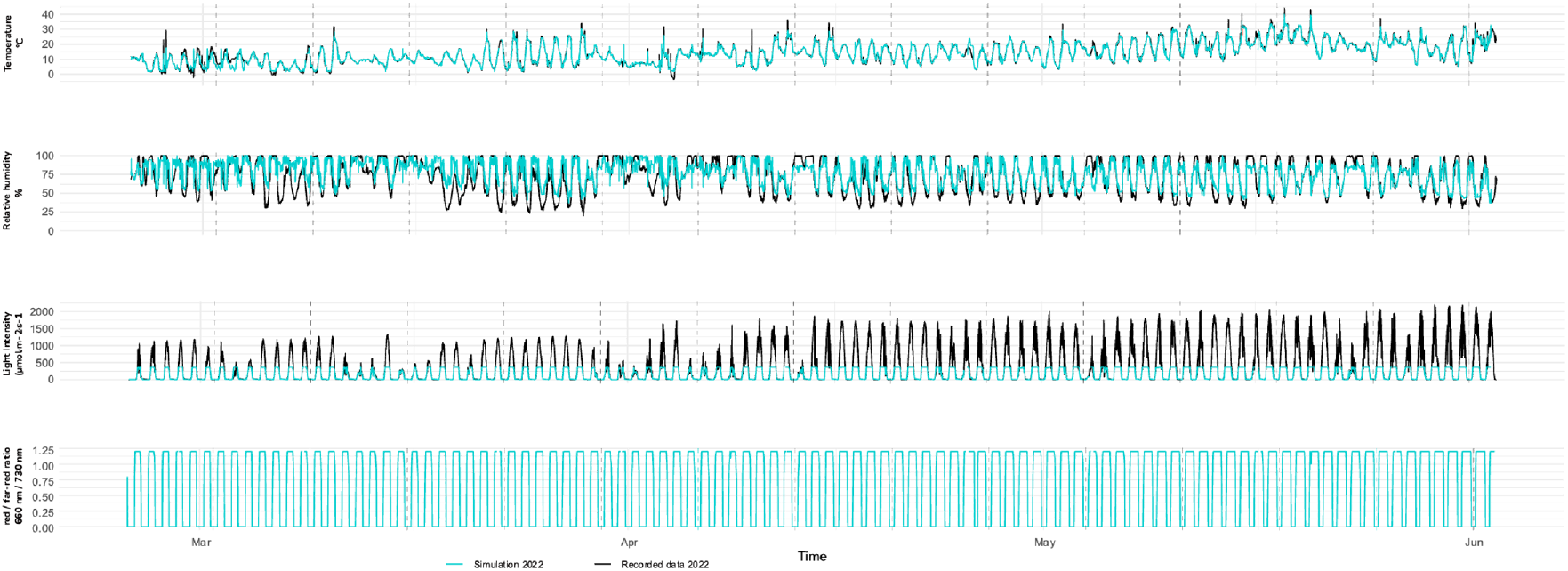
Environmental variables for the E&R experiment.

**Supplementary Figure 2.**
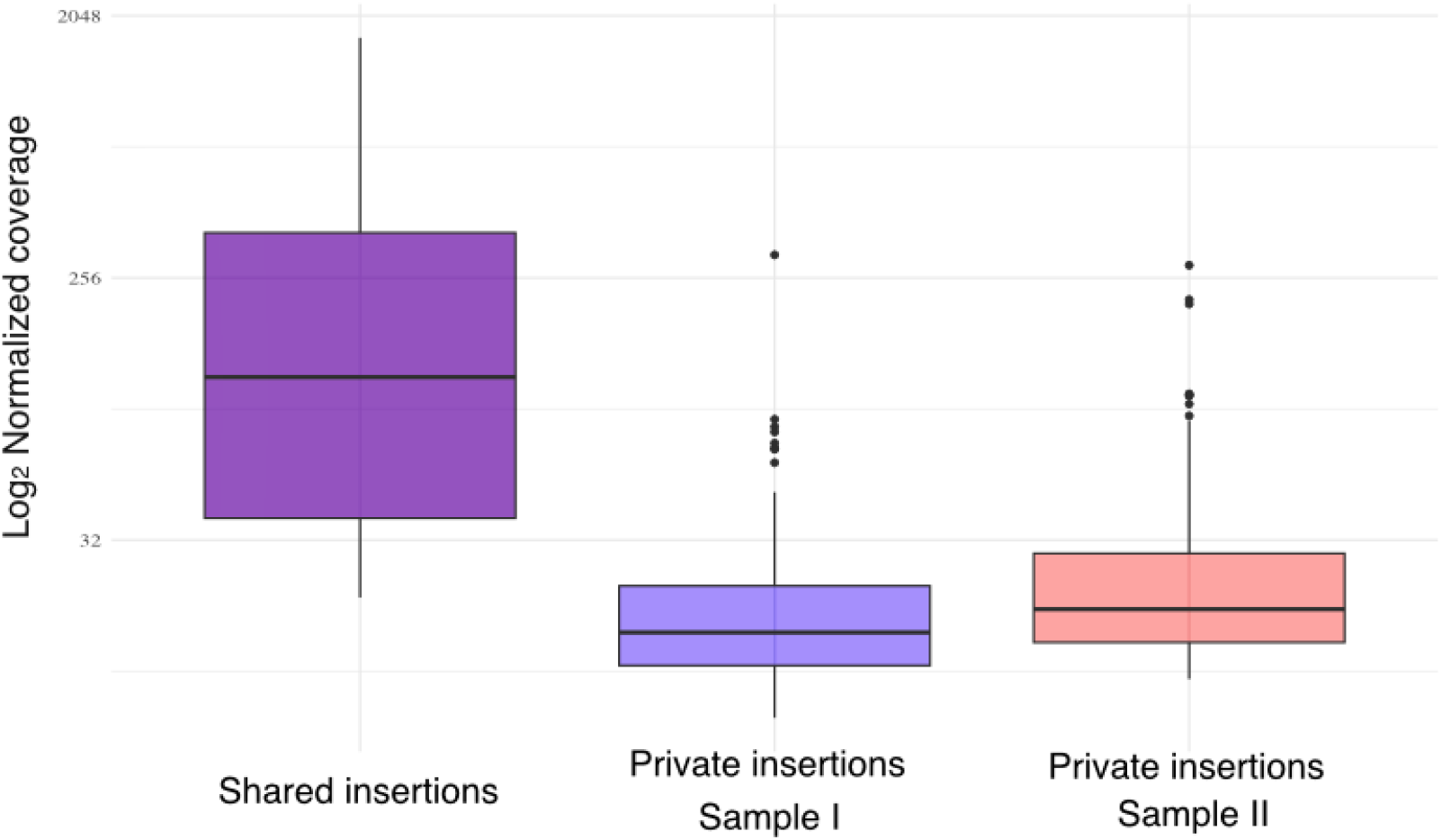
Number of TEd-seq reads supporting shared and private insertions

**Supplementary Figure 3.**
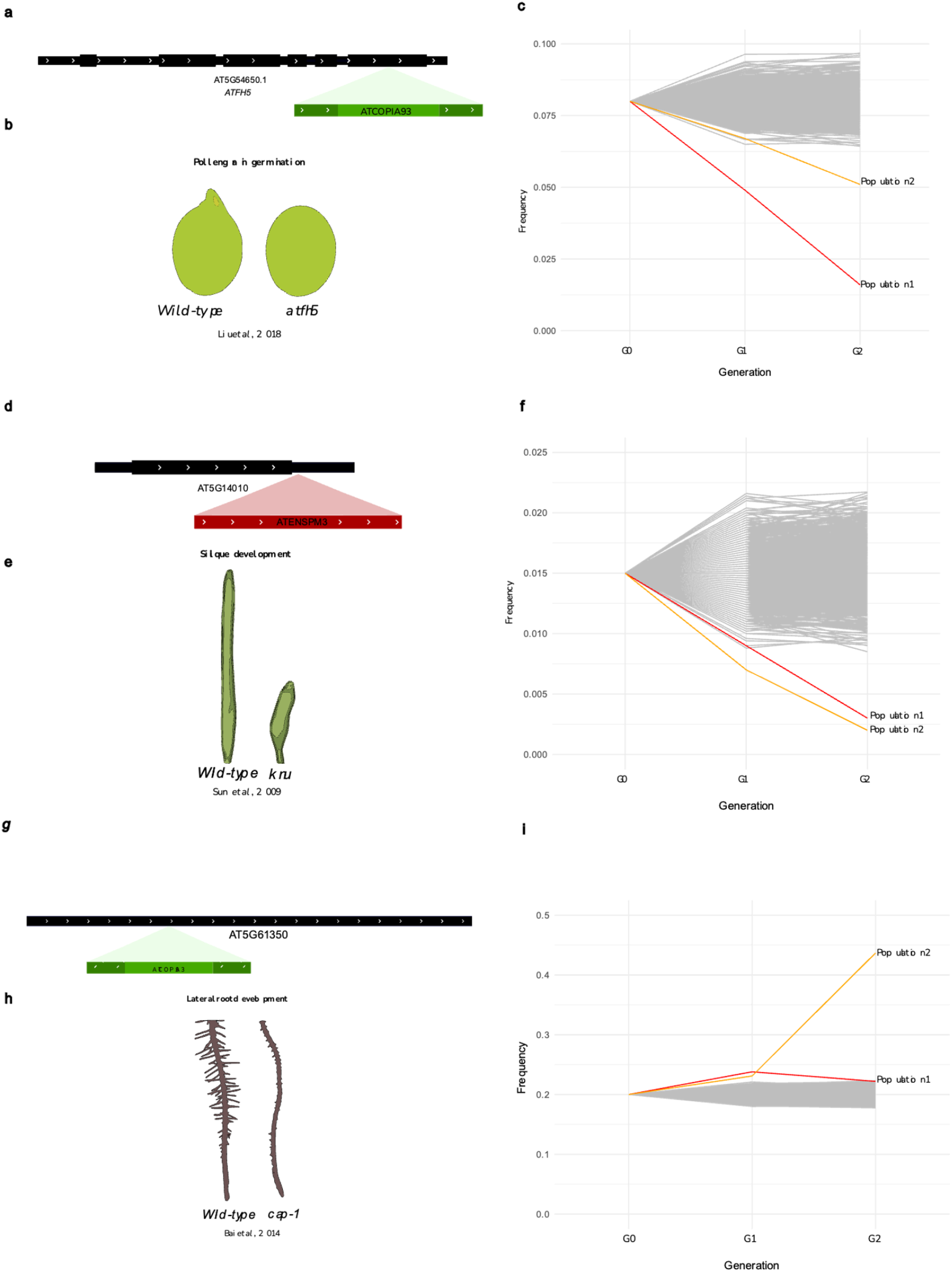
Examples of TE insertions showing significant changes in allele frequency across generations and their described phenotypic effects.

**Supplementary Table 1.**
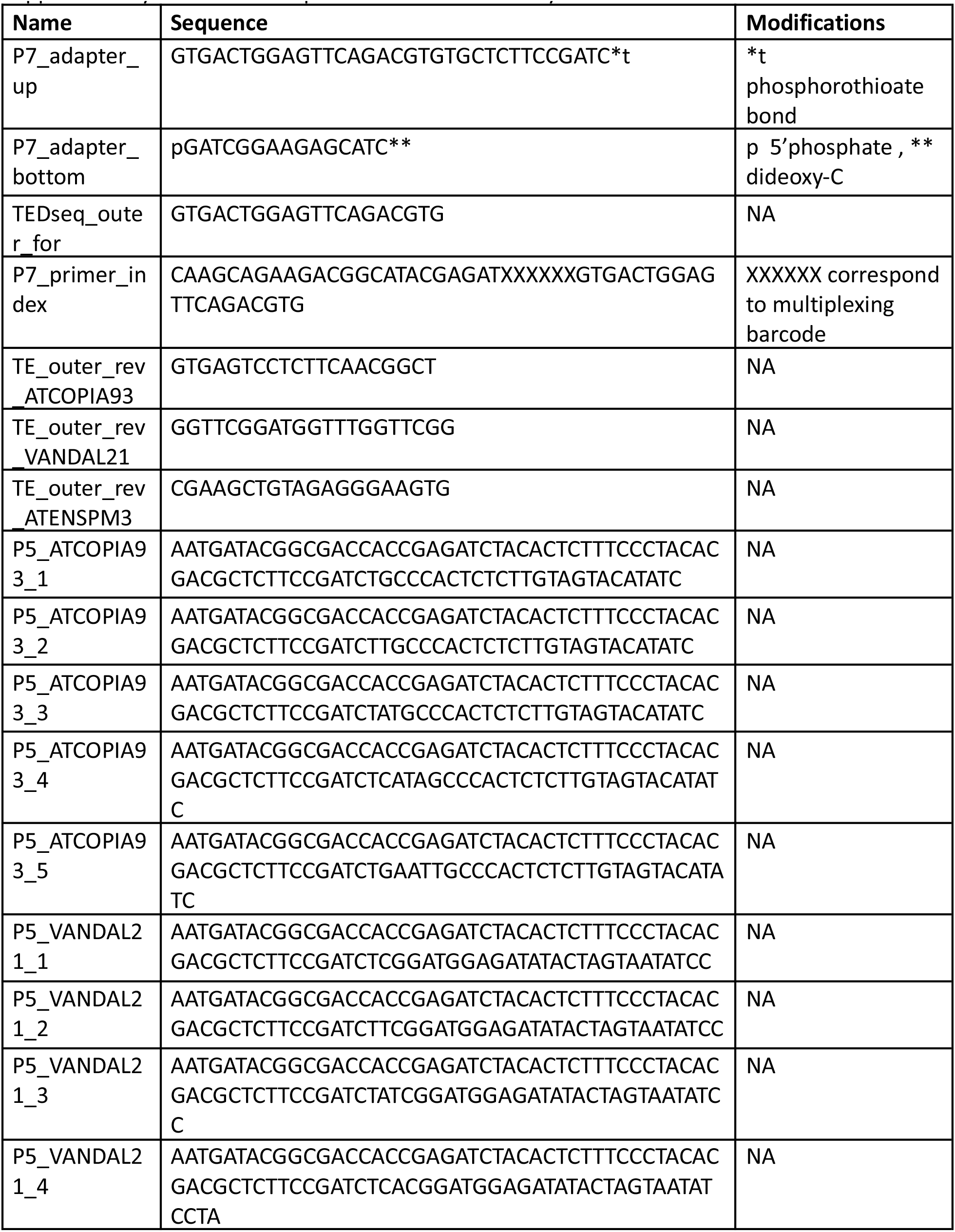
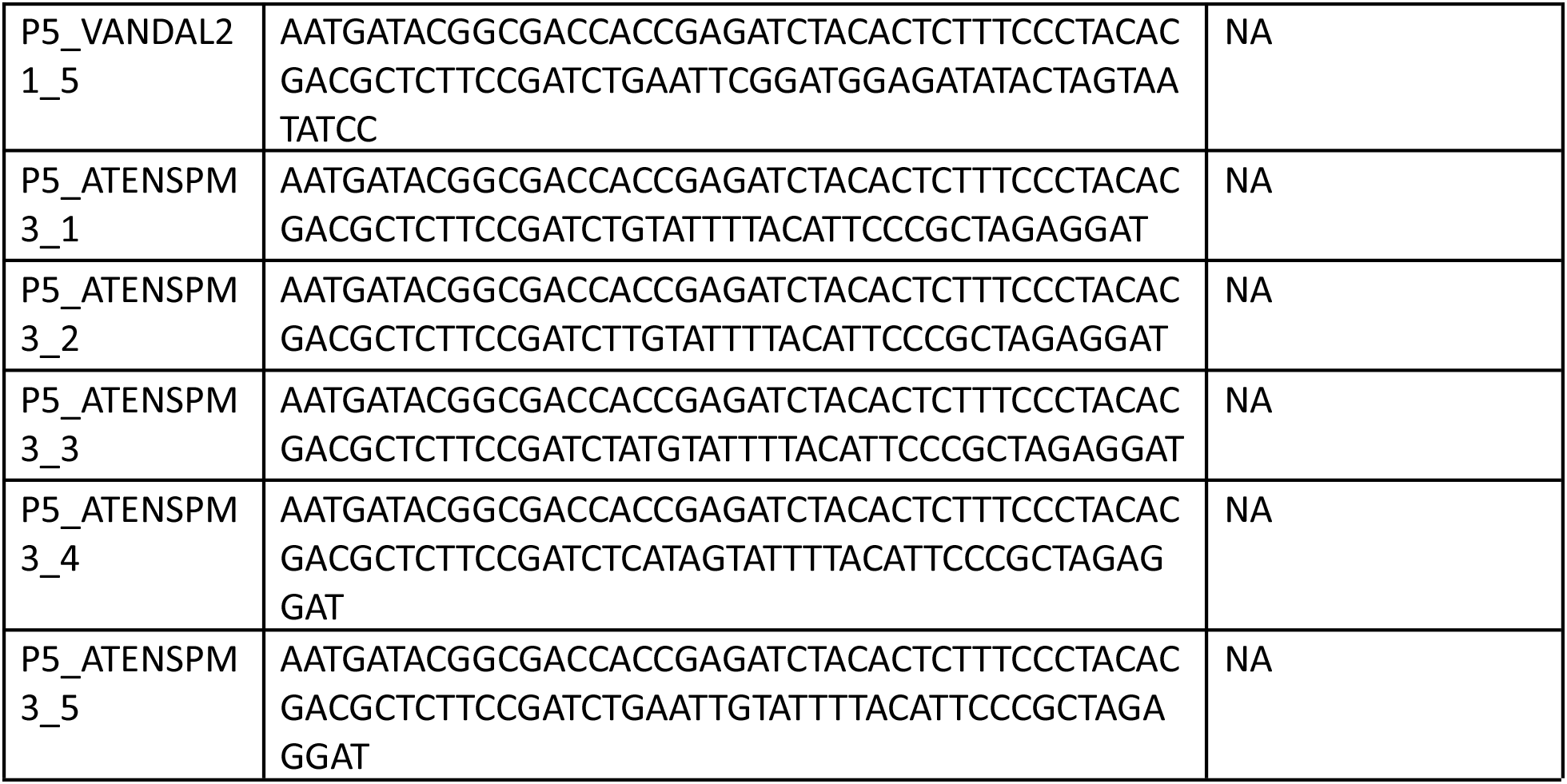
List of primers used in this study.

